# Changes in sea ice alter genetic structure of an iconic Arctic apex predator in less than three decades

**DOI:** 10.1101/2025.09.06.674276

**Authors:** Sean N. Vanderluit, Marie-Josée Fortin, Peter Van Coeverden de Groot, Rute B. G. Clemente-Carvalho, Evelyn L. Jensen, Andrea Gómez-Sánchez, Zhengxin Sun, Markus Dyck, Marsha Branigan, Stephen C. Lougheed

## Abstract

Climate change is having profound effects on biodiversity and species distributions worldwide. Nowhere are these effects potentially more pronounced than in the Arctic, where warming is almost two times the global average, and where year-round sea ice extent has significantly decreased, affecting many ice-dependent species. The polar bear (*Ursus maritimus*) is a circumpolar, apex Arctic predator, a sentinel of climate change, and a symbol of conservation. It is of immense cultural and spiritual importance to Inuit peoples and is hunted across the Arctic. Declines in sea ice have caused habitat fragmentation and loss, disrupting movement and prey access, potentially altering genetic structure and influencing polar bear’s potential to persist. Using samples collected from 1997 to 2020 by Inuit across much of the Canadian Arctic, 322 genome-wide autosomal DNA markers specifically designed to quantify polar bear genetic structure and a very stringent spatial-temporal method, we compare polar bear spatial genetic structure and landscape features between two periods across a consistent distribution:1997–2008 and 2009–2020. We observe marked changes in spatial genetic structure across the Arctic Archipelago over just two decades, shifting the boundaries between polar bear genetic clusters by ∼250 km. Landscape resistance models reveal the importance of sea ice and land cover type for each period, with spatial lag models showing that genetic change is best predicted by sea ice shifts between periods. Our study reveals rapid changes in genetic structure of polar bears in the Canadian Arctic, helps to inform conservation and management, and offers insight on future polar bear persistence across its immense, remote distribution.

## Introduction

The interplay among landscape heterogeneity, climate, life history, and micro-evolutionary processes shapes population structure, evolutionary trajectories, and geographical ranges, and ultimately whether species persist through time. Human activities are irrevocably altering the geographical distributions of species with many at increased risk of local or global extinction (Román-Palacios & Wiens 2020; Wiens 2016). For example, many species have undergone marked poleward or altitudinal range shifts in response to climate (Parmesan and Yohe 2003, Chen *et al*. 2011). Local populations of some taxa have also declined dramatically or disappeared altogether (Pounds *et al*. 2006, Atkinson *et al*. 2014, Ceballos *et al*. 2017, Tonina *et al*. 2022). Myriad species are now exposed to novel competitors (Alexander *et al*. 2015) and pathogens (VanWormer *et al*. 2019, Carlson *et al*. 2022) as range shifts cause geographic overlap. Responses to climate change vary with species ecology, including factors like dispersal propensity, diet, mating system, behavioral plasticity, and adaptive potential (Charmantier *et al*. 2008; Eckert *et al*. 2010; Travis *et al*. 2013; Razgour *et al*. 2019; Petherick *et al*. 2021). Different aspects of climate and geography also affect a species’ ability to persist, as shifting ranges may mean that species encounter thermal or precipitation maxima, or otherwise can be excluded by their climatic niche (Thomas *et al*. 2004; Pounds and Puschendorf 2004). The rapidity of environmental shifts lends urgency to the study of ecological change, and the associated implications for species distributions, ecosystem functioning, and evolution.

Compared to other regions, the Arctic is warming disproportionately quickly (Ratanen *et al*. 2022; Gauthier *et al*. 2024; Giesse *et al*. 2024), making it difficult to predict impacts on wildlife. However, the signatures of recent climate change are evident for many Arctic species, including significant degradation of the quality of breeding habitat of migratory birds (Wauchope *et al*. 2017) and the quality of caribou forage (Fauchald *et al*. 2017). Warming has also resulted in alarming declines in sea ice extent and volume, and changes in seasonality, with reported effects on Arctic biota that motivate detailed studies of mechanisms and interactions that underlie them (Post *et al*. 2013; Macias-Fauria and Post 2018; Kovacs *et al*. 2024). Earlier ice retreats have affected primary productivity with cascading effects throughout Arctic food webs (Leu *et al*. 2011), interrupted connectivity of large mammals (e.g. Peary caribou, *Rangifer tarandus pearyi*; Jenkins *et al*. 2016 and high Arctic Svalbard reindeer *R. tarandus platyrhynchus*; Peeters *et al*. 2019), and are predicted to cause diversity loss as more southerly species expand northward, bringing previously isolated species into contact (Kelly *et al*. 2010). These dynamics will be accentuated in the future, with increasing flux in ecosystems as ice-free conditions are expected before 2050 (IPCC 2021). Addressing these impacts requires research on how landscape changes impact ecosystems and populations, as well as community dynamics across northern latitudes. Examining how these changes will impact the genetic structure and distributions of keystone species in Arctic environments is crucial to understanding the fate of Arctic wildlife. However, comprehensive Arctic-wide studies remain challenging because of the remoteness of northern landscapes.

The polar bear (*Ursus maritimus*) is a circumpolar, apex Arctic predator and a symbol of climate change. In addition to its vital ecological role, the polar bear has cultural, spiritual and economic significance for Arctic Indigenous peoples – the Inuit. Globally, polar bears are delineated by 19 subpopulations (also called management units; Figure1a). Thirteen of these are entirely or partially within Canada, comprising two-thirds of the global population of between 22,000 and 31,000 individuals (Regehr *et al*. 2016). While western science projects imminent population decreases across the Arctic (Amstrup *et al*. 2008) and identifies declines in subpopulations at the southernmost extent of the range (SH, WH, FB – see Figure 1a; Canada’s Polar Bear Technical Committee 2018), Traditional Ecological Knowledge (TEK) often suggests stable or increasing numbers in some subpopulations (Stirling and Parkinson 2006; Tyrrell 2006; COSEWIC 2018). However, over most of its distribution, data on population trends are inadequate (Vongraven *et al*. 2018).

**FIGURE 1.**
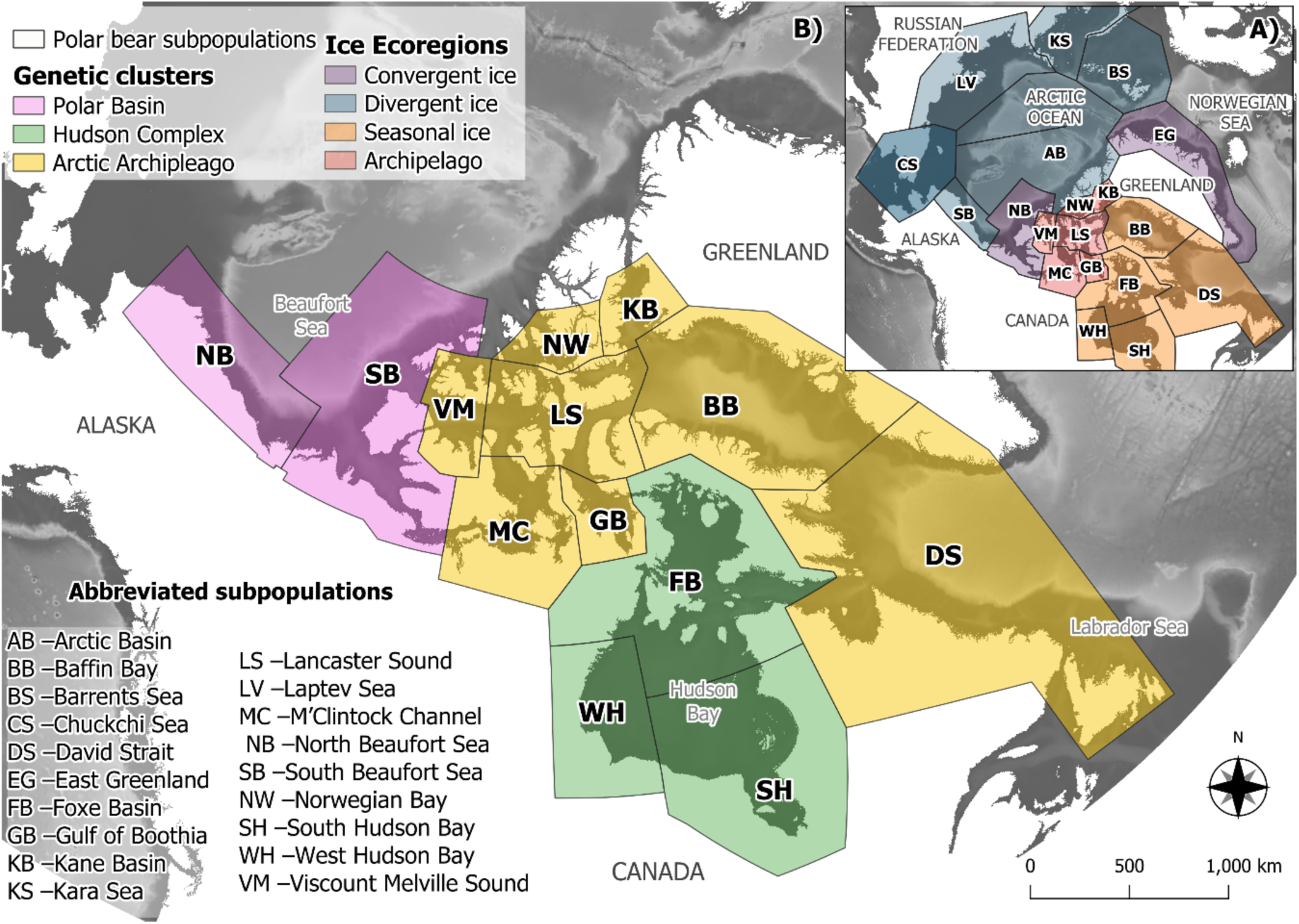
A) Global polar bear subpopulations that share seasonal patterns of ice motion and distribution (ice ecoregions) adapted from Amstrup *et al*. (2008). B) Subpopulations color-coded according to genetic clusters identified by individual majority assignment in Jensen *et al*. (2020).

Polar bear habitat is vulnerable because of expected increases in ice-free periods and summer ice loss (Hamilton and Derocher 2018). Situated at the northern extent of the Canadian Arctic Archipelago, the last ice area (LIA) is predicted to provide a last refuge for polar bears to persist up to the end of the 21^st^ century (Hamilton *et al*. 2014). The archipelago is also expected to have more transient bears as the century advances, sea ice deteriorates, and individual movements are shaped by adaptation/habituation to new landscapes, including more land-based and open-water travel. These effects may cause shifts in the polar bear distribution and concomitantly population genetic structure and the boundaries between genetic clusters (see below). Such shifts may be so profound that current subpopulation delineations upon which management is based will need to be revisited as landscape impediments to dispersal diminish in effectiveness and others arise.

Polar bears depend on several landscape characteristics for survival. Across the Arctic, bears use first-year or annual ice for hunting at the breathing holes of ringed seals (*Pusa hispida*), their primary prey. Ringed seals create and maintain breathing holes in annual ice, whereas thick multi-year ice precludes this (Smith and Hammill 1981). Thus, bears use sea ice to hunt and engage in breeding when sea ice is near its maximum extent, from April to mid-May (Lønø 1970, Rosing-Asvid *et al*. 2002; Stoops *et al*. 2012; Stirling *et al*. 2016). Bear distribution and population structure will be shaped by intra-annual dynamics. Sea depth of marine areas also influences the quality of polar bear habitat, with shallower continental seas promoting higher primary productivity and densities of ringed seals. For example, shallower areas are selected by polar bears in the Chukchi Sea between Russia and Alaska, despite high seasonal variation in distribution and availability of preferred habitat (Wilson *et al*. 2014).

Despite dependence on sea ice, regional differences in the landscape shape life history and phenology of polar bears exist. For example, sea ice seasonality influences the seasonal foraging and movement opportunities (Amstrup *et al*. 2008). In Hudson Bay, or the ‘Seasonal Ice Ecoregion’ (Figure 1a) virtually all sea ice melts and re-forms annually. In the Archipelago Ecoregion (Figure 1a), numerous fjords and persistent annual ice support high densities of ringed seals and thus polar bears. The convergent and divergent ice of the North and South Beaufort Seas in Canada (also ‘Polar Basin Ecoregion’; Figure 1a), is characterized by annual and multi-year ice and greater depths offshore with lower productivity. Studies of North American genetic structure have identified four broadly-distributed genetic clusters that largely coincide with these ecoregions: Polar Basin, Arctic Archipelago, Hudson Complex, and Norwegian Bay (Malenfant *et al*. 2016, Jensen *et al*. 2020; Figure 1b). Although these clusters are evident across datasets, individuals are often assigned to clusters hundreds of kilometers from their sampling location (Malenfant *et al*. 2016; Jensen *et al*. 2020), raising the possibility of movement over immense distances.

High genetic exchange among populations of polar bears is partly attributable to long-distance movements and vast home ranges (Ferguson *et al*. 1999; Johnson *et al*. 2017). Sea ice facilitates movement and foraging for polar bears, with evidence of genetic structure across hierarchical spatial scales (Campagna *et al*. 2013). Decrease in ice extent and duration is linked to changes in polar bear movement and constitutes the single most important predictor of their future abundance and distribution (Mauritzen *et al*. 2001; Parks *et al*. 2006). Loss of sea ice coverage has resulted in the loss of genetic diversity and disrupted gene flow in Barents Sea polar bears (Maduna *et al*. 2021). Telemetry-based movement studies document increasingly frequent long-distance swimming events that are physiologically costly responses to unsuitable sea ice coverage for dispersal and hunting (Durner *et al*. 2011; Pagano *et al*. 2012). Lower rates of survival and reproduction have also been linked to ice dynamics and coverage of continental waters in some populations (Regehr *et al*. 2010; Rode *et al*. 2010).

The rapid rate of decline in Arctic sea ice is already affecting Arctic biota, including polar bears, because of their dependence on this feature for foraging and movement. Rivkin *et al*. (2024) assessed the risk of climate maladaptation for Canadian polar bears using simulations and estimates of genetics offsets to forecast future environments. They suggest that subpopulations in the Canadian High Arctic (LS, NW) may be more vulnerable to climate change, and that ice thickness and temperature are among the most important predictors of allelic turnover. However, the genetic consequences of past sea ice loss across the Canadian Arctic on polar bears remain unknown. Here, we use genomic and spatial datasets spanning over two decades to investigate the effects of environmental changes, including sea ice extent, on genetic population structure across much of the Canadian Arctic. We (1) characterize broad-scale spatial genetic structure across Nunavut and the Inuvialuit Settlement Region, (2) assess landscape effects on genetic structure over marine and terrestrial environments, and (3) compare genetic changes over time. We predicted and indeed found that recent environmental changes, especially regional shifts in sea ice, have impacted genetic patterns over a surprisingly short timescale (just two decades), with shifts in polar bear spatial population structure across the Canadian Arctic.

## Materials and Methods

### Sampling, DNA extractions, library preparation and sequencing

Sampling, extractions, library preparation, and sequencing were a collaborative effort under the BEARWATCH project funded by Genome Canada. Samples of polar bear muscle tissue were collected by Inuit hunters during traditional harvests from 1997 to 2020 (permits ARI #WL 500540 to MB, WL-2018– 006 to SCL, WL-2019–061 to SCL). These dried or frozen muscle tissues were stored at -20°C or in 95% EtOH at -80°C, respectively, by the territorial governments of the Northwest Territories and Nunavut. Samples for BEARWATCH include biopsy tissue samples and harvest tissue ‘sets’ (comprising liver, fat, muscle, and colon). We subsampled harvest tissue samples with known provenance such that we had relatively even geographic and temporal coverage from various subpopulations, and approximately equal sample sizes of males and females. We extracted DNA using a salt extraction protocol adapted from Aljanabi and Martinez (1997) (S1), adding RNase A (Thermo Fisher Scientific Inc., Waltham, Massachusetts, USA) following lysis. We assessed DNA quality and purity on 1.5% agarose gels stained with RedSafeTM Nucleic Acid Staining Solution (iNtRON Biotechnology Inc., Seongnam, South Korea), and using a NanodropTM ND_1000 spectrophotometer (Thermo Fisher Scientific Inc., Waltham, Massachusetts, USA), and QFX Fluorometer high-sensitivity kit (DeNovix Inc., Wilmington, Delaware, USA), When necessary, we used the Faircloth and Glenn (2011) Serapure (GE Health Systems, Chicago, Illinois, USA) protocol for magnetic bead clean-up to remove fat and cellular debris (S2).

Double-digest restriction site-associated DNA sequencing (ddRADseq; Miller *et al*. 2007) libraries were generated following Peterson *et al*. (2012) and modified by Jensen *et al*. (2020) (S3). We also surveyed 322 genome-wide autosomal biallelic single-nucleotide polymorphisms (SNPs) and two sex-linked markers using Genotyping-in-Thousands by sequencing (GT-Seq) (Campbell *et al*. 2015), specifically designed to assay polar bear population structure following Hayward *et al*. (2021) (S4). GT-Seq allowed us to genotype samples with poorer-quality DNA, particularly older tissue samples that were unsuitable for ddRADseq.

ddRADseq libraries were sent to The Centre for Advanced Genomics (TCAG, Toronto, Ontario, Canada) for sequencing using two lanes of 125 base pair (bp) paired-end Illumina HiSeq 2500 in 2018 (Jensen *et al*. 2020). GT-Seq libraries were sequenced using Illumina MiSeq at Queen’s University, and Illumina NovaSeq 6000 at The Centre for Applied Genomics (TCAG).

### Genomic data demultiplexing and filtering

Genomic data were processed using a bioinformatics workflow developed for BEARWATCH (Jensen *et al*. 2020) (S5). Briefly, ddRADseq fastq files were demultiplexed based on P1 barcodes using process_radtags in STACKS v.2 (Catchen *et al*. 2013). GT-Seq fastq files were demultiplexed using i7 and i5 index sequence bases to separate plates and individuals, respectively, into fastq files using SAMtools v1.9 (Li *et al*. 2009). Reads were aligned to the polar bear reference genome (GCF_000687225.1; Liu *et al*. 2014) with BWA-MEM v0.7.17 (Li and Durbin 2009). Subsequently, we used BCFtools mpileup v1.13 to call variants of the targeted sites (Li 2011).

Using VCFtools v0.1.16, we filtered ddRAD to a minimum read depth of 6, and a genotype quality score of 18 (Danecek *et al*. 2011), then used BCFtools to combine files of individuals between batches. Using VCFtools, we ranked individuals by missingness to identify and exclude individuals with greater than 30% missingness, followed by 5% at the retained loci from further analysis. We then filtered individuals based on the useability of associated metadata and suitability for subsequent analyses with VCFtools. After filtering, 867 individuals were retained for further analysis (Figure 2a).

**FIGURE 2.**
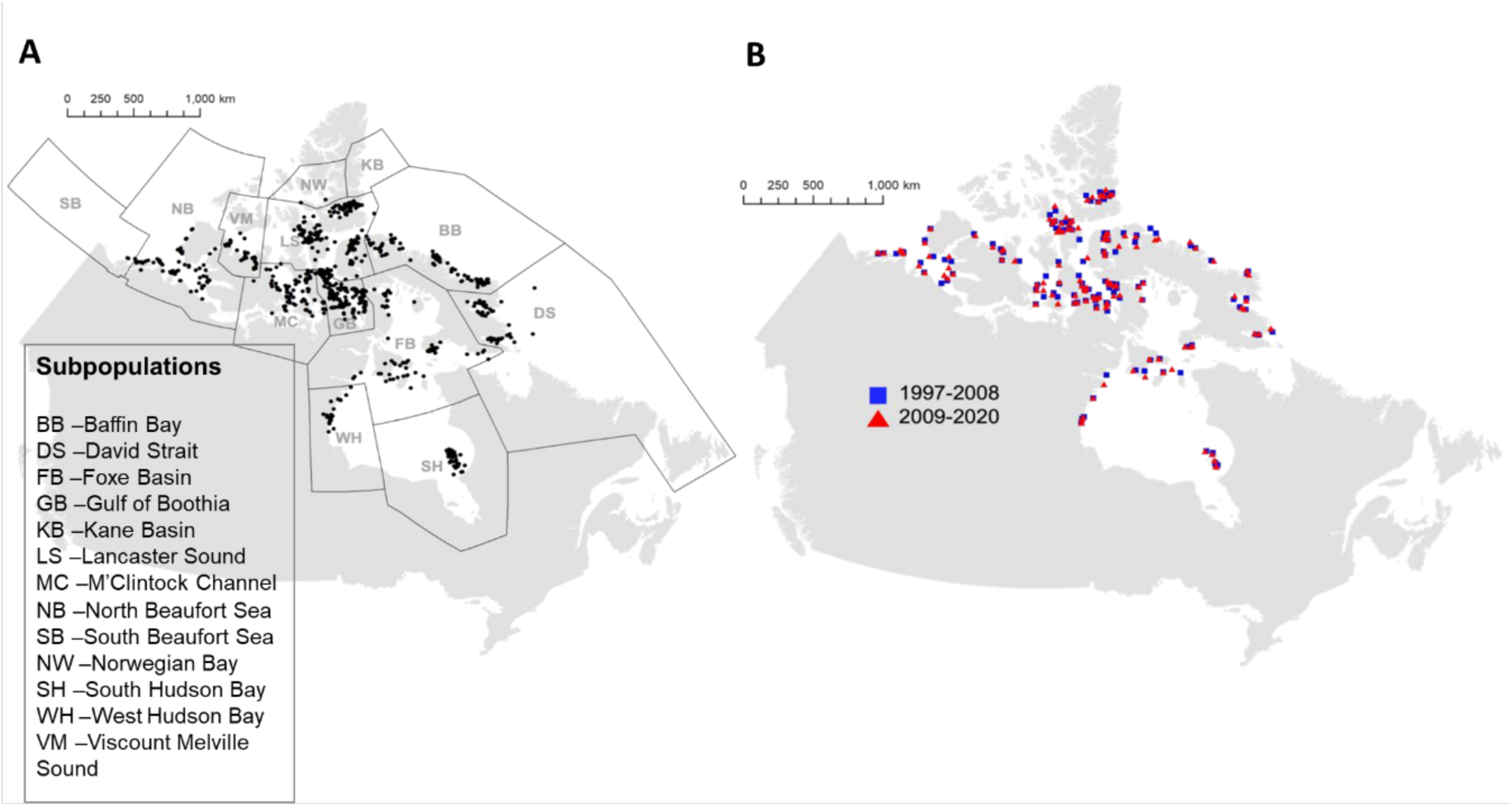
A) Distribution of 867 individual polar bear samples retained after filtering. B) Distributions of 126 individual polar bear sampling locations identified by areas of consistent sampling for 1997–2008 and 2009–2020 across Nunavut and the Inuvialuit Settlement Region.

### Areas of consistent sampling through time

We first apportioned the data into two periods to assess landscape genetic change: 1997–2008 (n= 421) and 2009–2020 (n= 446), representing roughly two polar bear generations (estimated ∼11.5 years – Regehr *et al*. 2016). As sampling was based on polar bear harvest quotas between 1997 and 2020, they may partially reflect hunter preference and community quota. To diminish spatial biases, we selected samples of acceptable quality using ‘areas of consistent sampling’ (Draheim *et al*. 2018) based on Voronoi polygons for the two distributions of samples using the ‘Create Thiessen Polygons Analysis’ tool in ArcMap 10.8. The resulting Voronoi polygons partition the overall sampling region into *n* cells, each centered on one of *n* sampled points; where every point within a cell is closer to the corresponding sample location than to samples in any other cell. The Voronoi polygons for the two periods were then overlain using the ‘Intersect analysis tool’ in ArcMap 10.8, identifying the areas that included at least one sample from either time-period. This stringent method allows us to delineate areas containing samples from both periods with diagnosed shifts attributable at least in part to local conditions that the individuals were exposed to (Draheim *et al*. 2018). We retained these samples for all subsequent spatial genetic analyses (n = 126 for 1997–2008 and n = 126 for 2009-2020, Figure 2b).

### Genetic and spatial genetic structure

In R v.4.1.1 (R Core Team 2021), we first performed a discriminant analysis of principal components (DAPC) for each time-period separately (n = 421 for 1997–2008 and n = 446 for 2009-2020) to depict broad patterns of genetic structure using existing subpopulation classifications, and used ‘optim.a.score()’ to select the number of principal components for PCAs with the *adegenet* package (Jombart 2008). We used the associated ‘find.clusters()’ function retaining all PCs to find K (the number of clusters) for each of these areas of consistent sampling.

To quantify the influence of straight-line distance on genetic structure for each time-period, we tested for positive spatial autocorrelation or isolation-by-distance (Wright 1943) using Mantel tests (Mantel 1967) among different established genetic clusters and simple Mantel correlograms in the R *ecodist* package with 1000 permutations (Goslee and Urban 2007). We calculated Euclidean genetic distances using this same package and geodesic geographic distances using the Geographic Distance Matrix Generator v1.2.3 (Ersts 2021), correcting for earth’s curvature.

We performed spatial analysis of principal components (sPCA) to assess spatial patterns of genetic variability for both periods using ‘spca()’ and ‘inverse distance connection networks’ in *adegenet* (Jombart *et al*. 2008; Jombart 2008). sPCA creates synthetic variables that maximize the product of entity scores and spatial autocorrelation (Moran’s *I*). As genotyped samples span Universal Transverse Mercator (UTM) zones 8 to 22, points were projected to EPSG:3978 (NAD83 / Canada Atlas Lambert). Missing data for each locus were replaced by mean allele frequency. Lagged principal component scores were interpolated and plotted to visualize genetic clines, and global and local structures were identified using ‘spca_randtest()’ (Montano and Jombart 2017).

### Landscape variables and Spatial analysis

We integrated land and sea cover data with pairwise genomic distances to evaluate the relative influence of landscape features on genetic structure. We retrieved historical ice extent data available weekly from 1997 to 2020 from the Canadian Ice Service (Environment and Climate Change Canada, Ottawa, ON, Canada). Further, we retrieved 2020 15 arc-second General Bathymetric Chart of the Oceans (GEBCO) topography and bathymetry data, and 30 m 2015 land cover data from the North American Land Change Monitoring System (NALCMS; Commission for Environmental Cooperation 2020).

We batch processed these data using the ‘Spatial Analyst Toolbox’ in ArcMap 10.8 (ESRI, Redlands, CA, USA). All ice extent data were retrieved in Canada Lambert Conformal Conic projection and converted to raster format. Raster data were reclassified into ice, land, or water, as these generalized classes offer the greatest consistency (Table S1). Raster data were mosaicked between Eastern, Western, and Hudson Bay regional extents per monthly observation. Embedded polygons with missing data were either reclassified to the likeliest adjacent cover type (e.g. small inlet classified as ‘NoData’ but enclosed by other ice types was reclassified to ‘ice’) or left as ‘NoData’ and excluded from cell statistics. These areas are peripheral to the core study area and inconsequential because we decreased the resolution of raster data (from 1 km grid cells to 10 km grid cells). Further, land extents were reclassified as ‘NoData’ before calculating cell statistics, and ‘NoData’ values were excluded from calculations to manage the effect of slight differences between individual monthly extents. A complete land cover raster was overlaid after cell statistics were calculated for each parameter.

Peak breeding occurs just after the annual ice maximum – for analysis, we used a global peak of April 22^nd^ based on the literature. Ice extents to reflect this are taken as close to this date as the CIS records that exist (ranging between April 17^th^ and May 1^st^). In the absence of a visually obvious pattern throughout time in these annual extents, 792 monthly extents for Hudson Bay, Eastern and Western regions were retrieved and processed to yield cell statistics across the sampling period (1997–2020); the 2020 data format was not compatible and was excluded. Monthly extents were taken for the earliest extent recorded for each of the three regions per month.

We clipped all raster data, buffering each individual retained from the areas of consistent sampling protocol, so that no individual was less than 600 km from a map edge, using ‘*Buffer and Extract by Mask*’ from the Spatial Analyst Toolbox in ArcMap 10.8 (ESRI, Redlands, CA, USA). We used this extent for all subsequent analyses to mitigate map edge biases (Koen *et al*. 2010).

### Resistance optimization

We parameterized resistance assignments of landscape variables using *ResistanceGA* v4.1-0.47 in R, rescaling environmental variable values based on a genetic algorithm within the ‘ga()’ function in the package (Scrucca 2013; Peterman *et al*. 2014; Peterman 2018; Shirk *et al*. 2018). Linear mixed-effects models were fit to the data using maximum likelihood population effects parameterization with Akaike Information Criteria (AIC) derived from the fitted model (Clarke *et al*. 2002). The algorithm subsequently selects the best values from the sample of randomly parameterized surfaces initially generated for a landscape variable based on the specified objective function. This retains the top 5% of models, and these undergo ‘mutation and crossover’. The entire process was repeated until 25 generations had passed without improvement in AIC. To reduce the number of cells, we coarsened landscape raster data to 10 km^2^ grid cells using the ‘Resample tool’ in ArcMap 10.8 (McRae *et al*. 2008; Peterman 2018).

Each time-period was optimized independently with 126 corresponding individuals and each resistance surface. *ResistanceGA* permits the optimization of either continuous or categorical surfaces (Peterman 2018). Polar bears occupy marine and terrestrial environments, complicating optimization of truly continuous terrestrial or marine surfaces. Thus, resistance optimizations were conducted on categorical surfaces with not more than 15 cover types per surface as a limitation of *ResistanceGA* (Peterman 2018). Each optimization was run at least twice to assess the convergence of parameter assignments (Peterman 2018). Single surface optimizations followed by a multi-surface optimization were done using ‘SS_optim()’ first, then using ‘all_comb()’ – a wrapper to ‘SS_optim(), MS_optim(), MLPE.lmm(), and Resist.boot(). all_comb()’ requires pairwise cost distances, implemented using the ‘gdistance’ library (van Etten 2017). For each optimization, a linear mixed effects model with maximum likelihood population effects parameterization (MLPE) was fit to the data. ‘Resist.boot()’ performs a bootstrap analysis with 1000 iterations, subsampling matrices of the response and distances created per surface, before refitting and calculating MLPE statistics.

### Spatial modelling of landscape and genetic change

We created genetic and landscape change surfaces based on data from the 1997–2008 and 2009–2020 areas of consistent sampling. We first created a landscape change raster based on the difference between averaged ice extents of either period using the ‘Spatial Analyst Toolbox’ in ArcMap 10.8. We then reclassified these based on the mean resistance assignment between separate single-surface optimizations from each period (Tables S2 & S3), generated previously using the ‘all.comb()’ function in *ResistanceGA* (Peterman 2018).

We created genetic change surfaces using the ‘Genetic Landscapes Toolbox’ (Vandergast *et al*. 2011), ‘Spatial Analyst’, and ‘3D Analyst toolboxes’ in ArcMap 10.8. The Single Species Genetic Divergence tool uses Inverse Distance Weighting and specified pairwise Euclidean genetic distances to weight links between sampling locations and to clip the area using a specified extent to avoid extrapolation beyond sampled areas. The result is a raster where each grid cell is assigned a weight based on the genetic distances among sampled locations (Vandergast *et al*. 2013). Using Draheim *et al*.’s (2018) framework, we subtracted the 2009–2020 standardized genetic landscape from the 1997–2008 landscape using the math algebra raster calculator from the ‘Spatial Analyst toolbox’. Positive and negative values in the resulting raster surface reflect local spatial genetic structure strengthened or weakened, respectively, between the two time periods. Landscape and genetic change surfaces were resampled to 25 km^2^ grid cells to manage computational constraints and, to account for spatial autocorrelation in the genetic response surface, analyzed using spatial simultaneous autoregressive lag models and spatial error models with the *spatialreg* package in R (Bivand *et al*. 2021).

## Results

### Broad-scale spatial genetic structure

The adegenet function ‘find.clusters()’ suggested a K of either 2 or 3 for both 1997–2008 (n = 421) and 2009–2020 (n = 446) for ddRADseq. Beaufort Sea, Hudson Bay, and Archipelago individuals form distinguishable clusters, with Archipelago individuals central to the other two (Figure 3a). This general pattern is also evident in the reduced representation dataset of 126 bears from either period of consistent sampling (Figure 3b).

**FIGURE 3.**
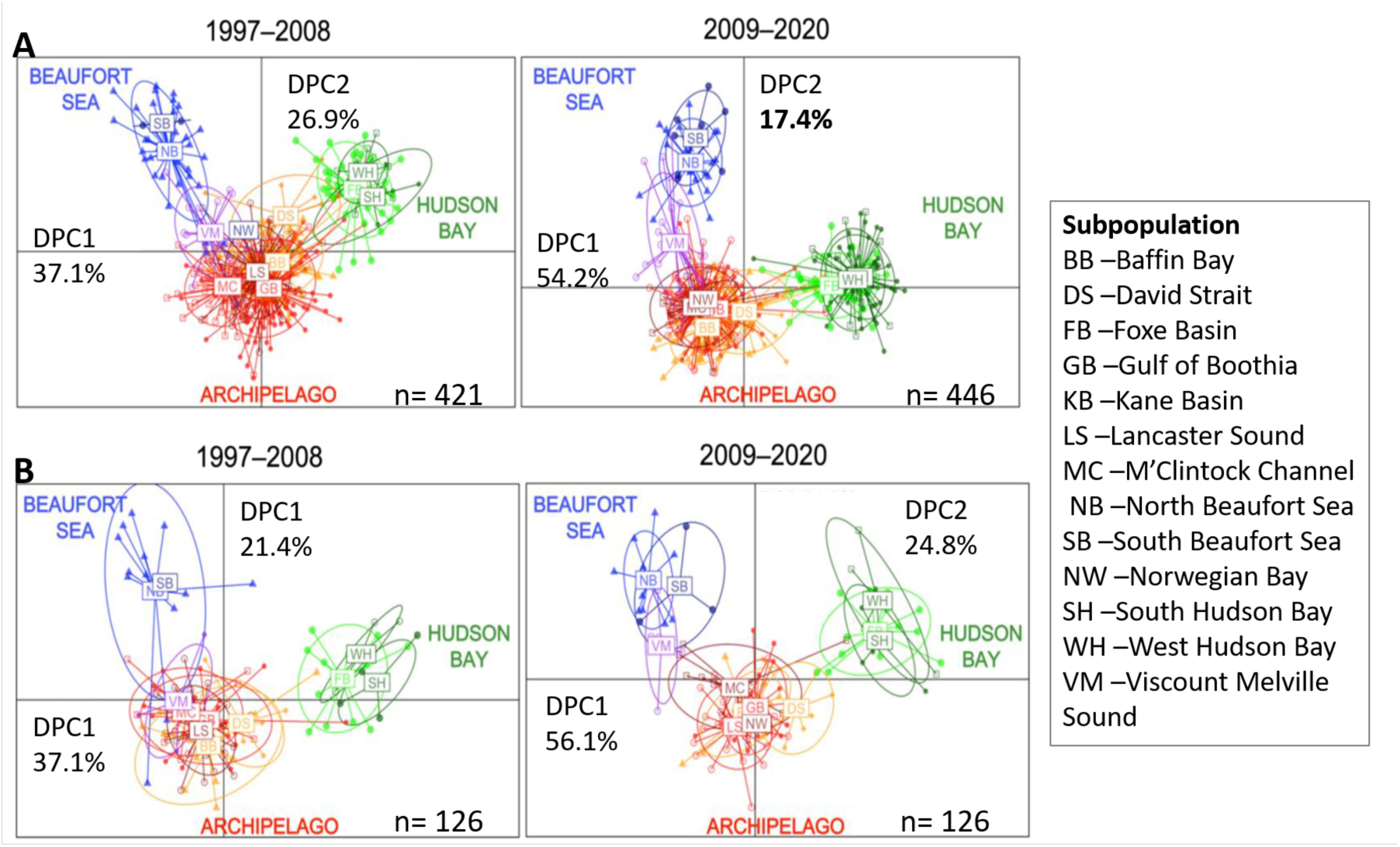
Discriminant analyses of principal components (DAPC) with 95% confidence ellipses around subpopulations for A) all polar bear samples for 1997–2008 (proportion of conserved variance=0.449, 51 PCs) and 2009–2020 (proportion of conserved variance= 0.483, 56 PCs); and B) areas of consistent sampling (1997–2008, proportion of conserved variance is 0.244, 12 PCs) and 2009–2020 (0.261,13 PCs).

**FIGURE 4.**
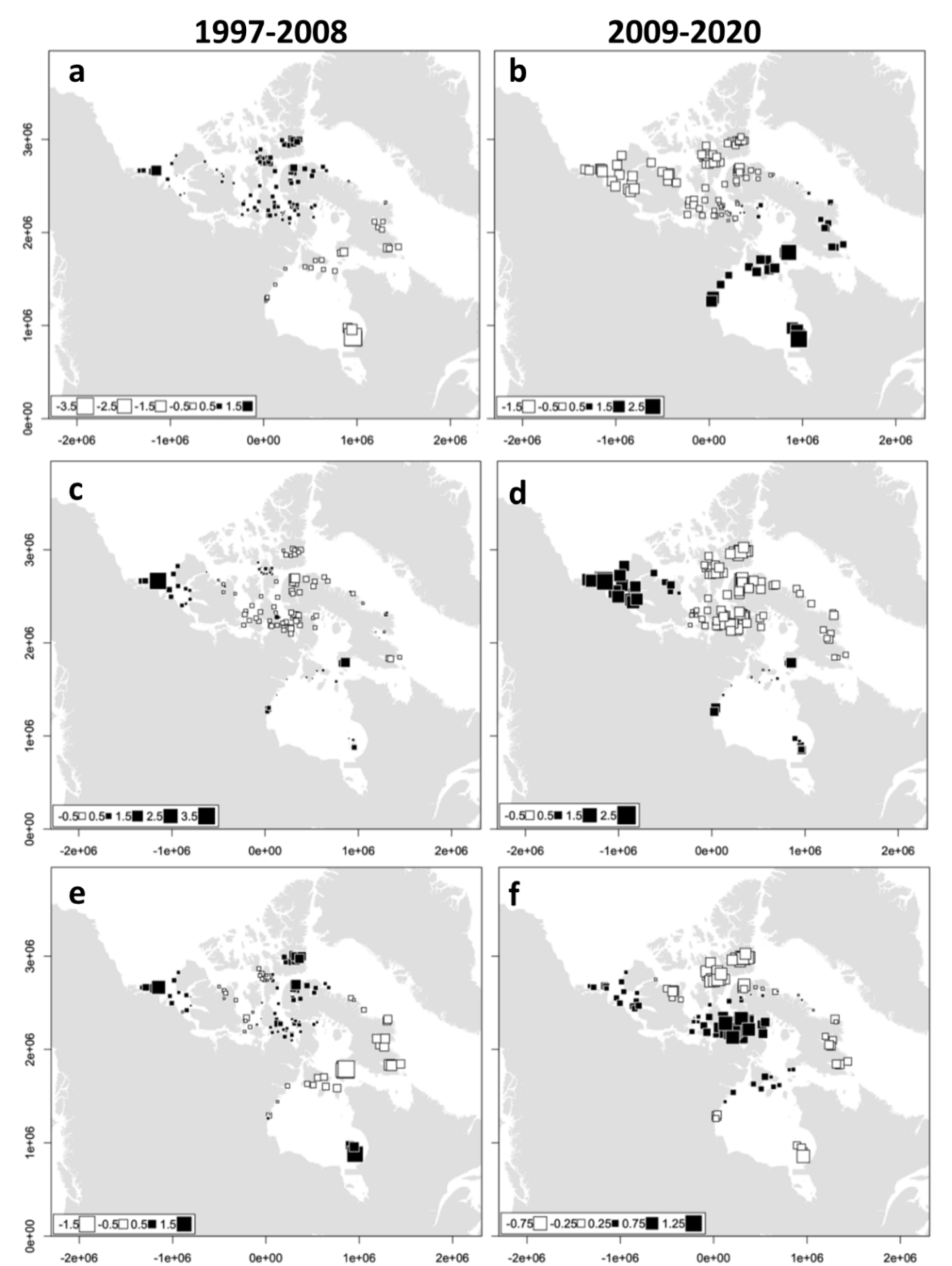
First (a-b), second (c-d) and third (e-f) axes of spatial principal component analyses for 1997–2008 (left, n = 126) and 2009–2020 (right, n = 126). Squares represent individual polar bear genotypes, with black or white indicating the sign, and larger squares indicating more extreme sPCA scores. Each is positioned by its sampling location across the Inuvialuit Settlement Region and Nunavut, Canada.

Among 126 consistently sampled areas, isolation-by-distance (IBD, Figure S1) is positive for 1997– 2008 (r_m = 0.32, p = 0.001), and for 2009–2020 (r_m = 0.24, p = 0.001). Mantel correlograms for 1997– 2008 (Figure S2) indicate positive spatial autocorrelation at the first four distance lags (Table S4). Beyond the fourth distance lag (> 856 km), sampled bears from this period are not more or less genetically dissimilar than random. The 2009–2020 Mantel correlogram shows significant positive spatial autocorrelation at the shortest three distance lags. Beyond 643.7 km, bears in this sample are not more or less genetically dissimilar than random (Table S4). These patterns are similar between individuals grouped by cluster, but significant IBD exists for Hudson Bay only in 1997–2008 and in the Archipelago for both periods (Table S5). Positive spatial autocorrelation among all samples partitioned by sex suggests a pattern of sex-biased dispersal for both periods, indicating greater mean dispersal among females (Table S6).

For each time-period, the sPCAs identify significant global structure (1997–2008: obs. = 4.47, P = 0.014; 2009–2020: obs. = 5.54, p = 0.002), but no local structure (1997–2008: obs. = 4.86, P = 0.296; 2009–2020: obs. = 4.42, p = 1.0). Significant global Monte-Carlo tests indicate the presence of positive spatial autocorrelation on at least one axis for each period (Table S7). The eigenvalue bar plots (Figure S3) and scree plots (Figure S4) indicate that the first three eigenvalues are appreciably larger than subsequent values and these were plotted for interpretation. The first global score distinguishes most individuals within the Hudson complex from the Archipelago and Polar basin bears for either time-period, with a cline in the Gulf of Boothia and Baffin Bay (Figure 5a-b). The second global score distinguishes Arctic Archipelago from both Beaufort and Hudson complex bears, with a cline in Viscount Melville (Figure 5c-d). The third global score depicts an east-west cline in the central Arctic Archipelago for the earlier period, and a north-south cline through the same area for the latter period (Figure 5e-f).

**FIGURE 5.**
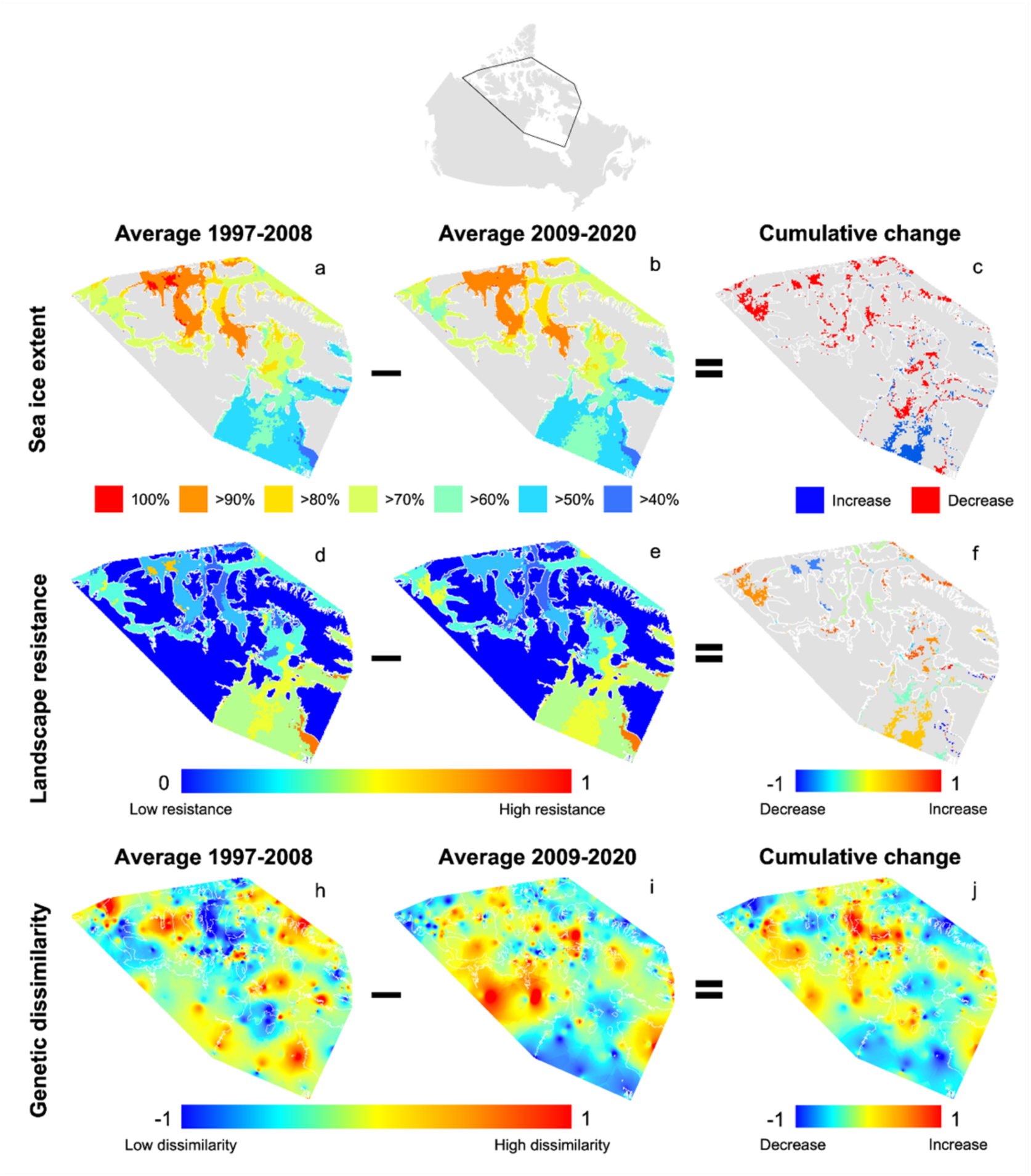
Sea ice extent, landscape resistance and genetic dissimilarity for 1997–2008 (a, d & g), 2009– 2020 (b, e & h) for polar bears across the Inuvialuit Settlement Region and Nunavut and cumulative changes over this period (c, f & i). Sea ice extent (10 km^2^) is based on retrieved and synthesized data from the Canadian Ice Service. Sea ice change between periods is classified by relative increase (blue) or decrease (red) between averaged extents across periods. Landscape resistance for both periods is based on ResistanceGA parameterized sea ice data (10 km^2^), standardized between -1 and 1. Landscape change was weighted to average ResistanceGA optimized resistance between periods. Genetic dissimilarity is based on Inverse Distance Weighted interpolations of pairwise Euclidean genetic distances among 126 consistently sampled areas (1 km^2^). Blue to red represents areas of low to high standardized genetic dissimilarity values ranging from 0 to 1. Positive values (red) indicate local spatial genetic structure strengthened, and negative values (blue) indicate local spatial genetic structure weakened, between periods. Coastlines are overlaid in white.

### Landscape resistance value assignment

Landscape resistances assigned from single surface optimizations of either period resulted in relatively low resistances, less than 25% of the greatest resistance assignment, to 80–90% and 90–99% ice extents for respective ice surfaces (Tables S2 & S3). For single surface optimizations among ice extents, land received the lowest assignment in 2009–2020, and second lowest in 1997–2008. For the land cover surface optimized by land cover class, marine received the lowest resistance assignment in 2009–2020 and second lowest in 1997–2008. Barren lands, the land cover type especially concentrated in the Arctic Archipelago, showed low resistance values (i.e. facilitate connectivity) for both periods.

For the 1997–2008 period, a simple distance model with uniform resistance across the study area was the highest performing model among all combinations of the average ice extent from 1997–2008 plus land cover, but was not significantly better than the null model representing no effect of distance (< 2 ΔAIC; Table 1). The associated bootstrap procedure identified the distance model as the top model across 1000 iterations (Table S8). For the 2009–2020 period, a combined land cover and sea ice model was the highest-performing model (Table 1), with the associated bootstrap procedure supporting this conclusion in 97.9% of 1000 iterations (Table S9).

**TABLE 1.** Linear mixed-effects models for 1997–2008 and 2009–2020 single and multi-surface combinations of ice and land cover, based on 126 sampled polar bear individuals from the Inuvialuit Settlement Region and Nunavut. Akaike Information Criterion values generated from the maximum likelihood population effects model, AICc-AIC value adjusted for the sample size, ΔAICc-difference between each model’s AICc and the lowest AICc. Weight indicates the average likelihood of the model.

| Surface | *k* | AIC | AICc | ΔAICc | weight |
| --- | --- | --- | --- | --- | --- |
| **1997–2008** |  |  |  |  |  |
| **geographic distance** | 2 | 19028.6 | 19028.6 | 0 | 0.60T |
| **null** | 1 | 19029.4 | 19029.4 | 0.79 | 0.40 |
| **ice1997_2008** | 13 | 19048.9 | 19052.1 | 23.50 | < 0.001 |
| **landcover** | 16 | 19055.5 | 19060.5 | 31.83 | < 0.001 |
| **landcover + ice1997_2008** | 28 | 19067.8 | 19084.5 | 55.90 | < 0.001 |
| **2009–2020** |  |  |  |  |  |
| **landcover + ice2009_2020** | 28 | 18719.8 | 18736.5 | 0 | 1 |
| **landcover** | 16 | 18803.4 | 18808.4 | 71.842 | < 0.001 |
| **distance** | 2 | 18871.7 | 18871.8 | 135.302 | < 0.001 |
| **ice2009_2020** | 13 | 19539.8 | 19543.0 | 806.498 | < 0.001 |
| **null** | 1 | 19671.2 | 19671.3 | 934.739 | < 0.001 |

### Landscape-genetic change

Genetic surfaces and genetic structure surfaces show local spatial genetic structure changed through time, with genetic dissimilarities accentuating in the Arctic Archipelago (Figure 5h-j). Although, landscape change is subtle compared to genetic change, there was a decrease in sea ice extent across the sampling region that resulted in changes in landscape resistance (Figure 5a-f). Spatial autoregressive lag models sampled at 25km^2^ showed genetic change is best predicted by a resistance change model, created by the difference between the best multi-surface optimizations for each period from the previous step (Table 2). Spatial error models at 25 km^2^ also identified a resistance change model as the best performing (Table S10).

**TABLE 2.** Spatial autoregressive lag models to predict polar bear genetic change (GC) across the Inuvialuit Settlement Region and Nunavut using two metrics of landscape change. RES – change in optimized resistance between periods, averaged between sea ice parameterizations for either period DEG – direction of sea ice change in either positive or negative degrees of change. wAIC is the average likelihood of the model.

| **Model (GC ~)** | **$\rho$** | ***P*** | **AIC** | **ΔAIC** | **wAIC** |
| --- | --- | --- | --- | --- | --- |
| βRES | 0.996 | < 0.001 | 50392 | 0 | 0.62 |
| βDEG + βRES | 0.996 | < 0.001 | 50393 | 1 | 0.37 |
| βDEG | 0.996 | < 0.001 | 50401 | 9 | 0.01 |

## Discussion

Rapid environmental change can have dramatic effects on biodiversity, causing shifts in distributions and population declines. Delineating specific landscape changes that influence species distributions, dispersal, and genetic structure is crucial for predicting how species will respond to environmental change across their geographic ranges. Even over small timescales, changes in landscape and climate may exert important effects on intraspecific genetic diversity (Draheim *et al*. 2018). Here, we combined over two decades of polar bear genetic data from 322 independent, genome-wide SNPs specifically designed to capture polar bear genetic structure across the vast Canadian Arctic with geographic land and sea ice cover data, and performed a stringent spatial-temporal landscape-genetic comparison. We observed changes in genetic structure that reflect altered seasonal sea ice conditions across Nunavut and the Inuvialuit Settlement Region. Bears sampled across equivalent spatial distributions in the 1997–2008 and 2009–2020 periods of consistent sampling showed marked changes in spatial genetic structure. A decrease in average sea ice extent between periods correlated with this genetic change. This implies that climate change-driven diminutions in sea ice have already had tangible effects on polar bear population structure over this very short time span.

Polar bears rely on annual or first-year ice for foraging and are forced onto multi-year ice or land each year when these habitats disappear. These annual movements are dictated by seasonal ice break-up and formation in Hudson Bay (Parks *et al*. 2006; Biddlecombe *et al*. 2021) and in the Beaufort Sea (Amstrup *et al*. 2000; Johnson and Derocher 2020; Pagano *et al*. 2021). Across these regions, swaths of multi-year ice have been replaced by annual ice and annual ice itself has been replaced by longer ice-free periods; these patterns are evident in the averaged extents over the last two decades. While shifts from multi-year to annual ice may increase marine productivity because of greater light penetration, result in higher seal densities, and augment polar bear foraging opportunities at higher latitudes, contractions of annual extents at lower latitudes negatively impact polar bears at these limits of their distribution. Bears in the regions undergoing rapid environmental change may have limited opportunity to forage and suboptimal foraging habitat due to ice movement and distribution, or may encounter physical impediments to movement between optimal foraging platforms, as swimming for these bears is energetically costly (Griffen 2018).

### Spatial genetic structure

Patterns evident from temporal comparisons using a suite of analytical approaches (DAPC, sPCA, and mixed effects models) from each of 126 spatially consistent sampling locations indicate differences in the genetic structure between the 1997–2008 and 2009–2020 sampling periods. Overall, genetic structure increased over time, and subpopulations at the peripheries of ecoregions experienced relatively more genetic change. Despite shared environmental dynamics of the Seasonal Ice Ecoregion that includes West Hudson (WH), South Hudson (SH), and Foxe Basin (FB) of the lower Hudson Bay complex (Amstrup *et al*. 2008), Davis Strait (DS) and Baffin Bay (BB) clearly cluster with individuals from Arctic Archipelago subpopulations including Gulf of Boothia (GB), Lancaster Sound (LS), M’Clintock Channel (MC) and Norwegian Bay (NW) in both sampling periods (Malenfant *et al*. 2016 and Jensen *et al*. 2020). Genetic continuity that we find across higher latitudes may bode well for the persistence of these subpopulations, although they may face higher risk of maladaptation to climate change (Rivkin *et al*. 2024). Clustered samples from SH and WH are each genetically distinct from subpopulations in the rest of the distribution. However, these gaps are at least partially a consequence of long ice-free periods in much of Hudson Bay, which restrict polar bears from foraging in these areas for almost half of the year. Nevertheless, FB further to the north clusters closely with these lower Hudson subpopulations, and only diverges on the third axis of sPCAs from South Hudson, implying that bears do not readily migrate between the Hudson complex and Archipelago.

The spatial genetic disjunction (sPCA 3^rd^ axis) between SH and WH likely reflects a history of isolation, attributable to ice dynamics in Hudson Bay. This disjunction may, in part, be caused by the directional nature of annual ice; freeze-up from the northwest to the southeast, and break-up southward from the coastal fringes of Hudson Bay (Danielson 1971). These genetic differences may also be accentuated by increasingly shorter ice periods (October–August) over this extent trending towards earlier break-ups in the southern and western parts of Hudson Bay, and later freeze-up in the north and northeast (Gagnon and Gough 2005). The advance of this earlier pattern of break-up and later freeze-up in tandem with fidelity to summer areas may effectively isolate SH bears, which otherwise lack meaningful connectivity to the remainder of the polar bear distribution (Stirling *et al*. 2004).

Viscount Melville (VM), and to a lesser extent MC, appear to cluster closer to Beaufort subpopulations of the polar basin (South Beaufort (SB) and North Beaufort (NB)) in the more recent period (2009–2020), with the boundary between genetic clusters shifting by 250 km. This is probably the result of dispersal from the Beaufort region – shifts in where the bears of this region now occur – driven by loss of multiyear ice from the Beaufort Gyre and parts of the Archipelago, and as intra-annual extents retreat from the mainland for longer periods of the year (Fischbach *et al*. 2007; Gleason and Rode 2009; Kwok and Cunningham 2010). This implies increase in preferred habitat at higher latitudes and eastward towards the Archipelago with shallower seas and high intra-annual ice, and decrease in preferred habitat at southern latitudes.

### Landscape-genetic resistance and change

Comparing resistance models optimized and ranked for each period, greater resistance and landscape effects were observed in the later period. The 1997–2008 period showed no discernable resistance attributable to landscape implying that physical distance was the overarching factor shaping genetic structure, consistent with other studies (Rivkin *et al*. 2024). In contrast, the top model for the 2009–2020 sample was a combined average ice extent and land cover surface model. This implies decreasing connectivity and greater influence of landscape through time. Various forest types, shrubland, and grassland all converged on relatively higher resistance assignments based on genetic distances and are not habitats typical of polar bears. Snow and ice cover occurs primarily at high latitude, and higher elevation regions of the Archipelago (e.g. eastern Baffin Island and Devon Island); thus, high resistance may reflect the added barrier of higher elevations. Barren land cover types comprise the least resistant category, possibly due to the distribution of these areas on Archipelago islands, where higher densities of bears occur. Our optimized resistance assignments for ice extents are also supported by other sea ice selection models for polar bears (Durner *et al*. 2009; Lone *et al*. 2018), where polar bears select intermediate to high ice concentrations where they would have greater hunting success; thus, it is reasonable that these surfaces would represent lower relative resistance to dispersal. However, resistance models can misrepresent true landscape-genetic relationships (Draheim *et al*. 2021), and it would be fruitful to further ground-truth these landscape resistance models using individual movement data (Cushman and Lewis 2010). Although the addition of topographic and bathymetric variables as well as greater specificity among categorical land and seascape variables may improve model performance (Draheim *et al*. 2021), our results are clear that sea ice concentrations and land cover composition influenced polar bear genetic structure between these time periods.

Genetic change was localized in some areas of the Arctic. Local genetic structure in parts of the central Archipelago increased throughout time, which was surprising as this region has historically had relatively constant and favorable conditions (Barber and Iacozza 2004). Of these, the most pronounced changes appear to be in a region through the GB that dramatically increased in local genetic structure between periods. This area has a shorter ice season, with coverage reduced by 10% between the two time periods. Slight changes in the availability of ice, or rather the seasonal dynamics of the interceding years, may have driven this shift in structure, as ice declined east of the GB and on the west coast of the Boothia Peninsula. These regional dynamics and incipient isolation are relevant to the continuity between GB and LS, and between MC and GB. These subpopulations comprise the core of the central Archipelago region, the highest densities of bears globally, and perhaps the greatest promise for the persistence of this species in the Arctic’s Last Ice Area (LIA) (Newton *et al*. 2021). These changes could be due to the wide geographic dispersal of individual bears from more distant and genetically disjunct regions, such as the Hudson Complex and Beaufort Sea. Decline in the ice period, occurring at greater rates in these regions (Gagnon and Gough 2005; Fischbach *et al*. 2007), and the Barents Sea and Arctic Basin (Stern and Laidre 2016), may be driving movement at local scales, as bears converge on areas with relatively longer ice periods to hunt. As temperature continues to increase, this will continue to reduce sea ice across the Canadian Arctic (Derksen *et al*. 2019). This means that we may see increased emigration from these regions as seasonal ice permits or restricts individual movements, decreasing global spatial genetic structure over the long term, but increasing local genetic structure in the interim. These shifts of individuals may be beneficial if some of these subpopulations possess local adaptations to higher temperature or longer ice-free periods, offering a possibility for persistence in new conditions (see Rivkin *et al*. 2024). Nonetheless, regions experiencing the greatest relative changes in intra-annual ice coverage will have the greatest risk of local extinction.

Decreases in local genetic structure are evident outside the Archipelago. We found a decrease in structure between GB and MC through time across the genetic change surface and in the third axis of the 2009–2020 sPCA. This is contrary to findings of a previously undiscovered cluster (Hayward *et al*. 2021), although the authors note the possibility of a temporal confound being the cause of this MC cluster. This shift in genetic structure, irrespective of the underlying cause, exemplifies the rapid shifts in species population structure that may be revealed by temporal studies like ours, and cautions against combining data from samples collected over multiple generations. Further, decreases in genetic structure throughout much of the Hudson Complex, Beaufort, and along the north coast of Baffin Island may signify significant shifts caused by dispersal between time periods within a single polar bear generation. These dynamics likely influenced lower correlation between inter-individual genetic and geographic distances over consistently sampled areas of either period, as well as within each of the three main clusters. This implies increasing dispersal over the study area perhaps due to later annual sea ice freeze-ups and earlier break-ups across much of Hudson Bay, the Archipelago, and Beaufort Sea. These effects will likely become more pronounced as polar bear habitats continue to contract across the Canadian Arctic.

The difference in amount of ice and the relative change in resistance, and a spatial lag term predicted genetic change between periods in our spatial regression models. Although landscape change appears relatively small-scale within the bounding area used in our analyses, more dramatic shifts in ice coverage occurred adjacent to this area with impacts on associated population structure in the lower Hudson Bay, offshore of Baffin Island, and in the Beaufort Sea. The inferences made based on the spatial regressions imply a general causal relationship between genetic and landscape/seascape change throughout these diverse ecoregions of the Canadian Arctic.

### Caveats

The value of resistance models to infer polar bear movement depends on whether the samples we used accurately capture the spatial and temporal dynamics within the study region. Historical gene flow and processes from decades prior to the present study period may have contributed to these genetic effects, as genes may move over greater distances over multiple generations than any single individual (Bohonak 1999). Many of these inferences also rely on the assertion of rapid genetic change between two consecutive 12-year periods, but this is supported by results from a simulation study showing that wide-ranging taxa with high propensity for dispersal can exhibit individual-based genetic signatures of barrier detection if landscape connectivity changes abruptly (Landguth *et al*. 2010). Shifts in genetic structure may result from dispersal from outside our study area, a proposition that we were unable to evaluate as we lack sufficient samples from some contiguous subpopulations including Norwegian Bay (NW), Kane Basin (KB), Chukchi Sea (CS), and Arctic Basin (AB). However, we note that our sample coverage spans a significant portion of the extant polar bear distribution. Sex-biased dispersal was evident, albeit subtle, with philopatric females showing a greater correlation with pairwise distances, implying less dispersal, than males (Tables S4 and S6); however, these differences may be due to the movement limitations of females accompanied by cubs (Amstrup *et al*. 2001; Taylor *et al*. 2001). While selecting for areas of consistent sampling ensures a temporally representative sample across space (Draheim *et al*. 2018), there is still some clustering across the sampling area (Figure 2b), which may impact interpretations (Oyler-McCance *et al*. 2013). As the available samples are harvested individuals, we could not use a different sampling regime without severely reducing the samples for temporal analysis; thus, samples are clustered presumably where both densities and quotas are high near communities, and hunting access is permitted based on seasonal conditions. Despite these caveats, we detected a significant link between landscape and spatial genetic structure change over a remarkable short time span.

### Implications

Humans, particularly via the wildlife management programs of the 20^th^ century, may have an underappreciated effect on historical and existing gene flow and population structure of polar bears. For example, the M’Clintock Channel subpopulation has been under a moratorium on harvesting since 2000 due to suggested overharvesting between 1970–2000 (US Fish and Wildlife Service 2001; Taylor *et al*. 2006). These restrictions were imposed after dramatically conflicting evaluations of population status between 1970s and 2000s; the legacy of this moratorium and the uncertainty of population size preceding warrant additional study especially given the profoundly negative impacts on the local Inuit community (de Wildt 2022). Prior to the 1970s, declines in polar bears attributed to spikes in recorded harvests and unknown numbers of unrecorded harvests throughout the preceding decades lead to the 1973 signing of the International Agreement on the Conservation of Polar Bears; this coincided with a rapid rise in human population growth within the Arctic Circle (Prestrud and Stirling 1994; Atwood and Wilder 2021). The legacy of these purported historical overharvests may continue to influence contemporary spatial genetic structure.

Maintaining connectivity for climate-induced habitat shifts is an important conservation goal (Tiberti *et al*. 2021). Though the scale of these efforts varies greatly among species differing in life histories and distributions, the results can inform directed actions, hunting or industry regulations, or delineation of coastal and marine protected areas between disjunct occupied portions of a species’ range. Arctic commercial activities are expected to rise as access increases to previously inaccessible areas for shipping routes and resource exploration, and various states attempt to assert ownership over these resources (Carnaghan and Goody 2006; Huntington et al. 2023). These activities incur an additional dimension of risk for the future of polar bears and their persistence. Conflicts with other bear species (grizzly and black) as well as humans will also invariably increase as annual ice extents continue to shrink, and energetically-stressed bears spend more time away from ice and potentially nearer coastal communities (Stirling and Parkinson 2006; Towns *et al*. 2009). Continued robust and science-based management (Regehr *et al*. 2017), based on modelled sea ice extents under climate scenarios, will help to identify and preserve refugial polar bear habitat and manage rising instances of human-bear interactions.

To our knowledge, ours is the first to study directly temporal changes in polar bear genetic population structure. Most polar bear research is done in isolated studies examining the effects of climate change on life history and ecology or describing population genetic structure in a single temporal sample (Vongraven *et al*. 2018). Over a distribution spanning the Arctic Archipelago, we detected short-term genetic change and increasing isolation effects of the landscape, which may confound evaluations of population structure made using genetic samples pooled from harvests conducted over many years. Land cover and averaged sea ice extent influenced polar bear genetic structure in the latter sampling period, and the change in ice extent between periods appear to have caused genetic change over just over two decades. Further research on the Arctic Archipelago populations is needed, but not without the consideration of Inuit knowledge from the outset as a valuable source of insight or consultation with Inuit communities. Interdisciplinary and temporal studies such as ours can fill important knowledge gaps and reveal previously overlooked effects that afford additional insight into evolutionary processes and strengthen directed action to conserve biodiversity (Stange *et al*. 2021), especially important in these remote, vulnerable Arctic landscapes.

## Acknowledgments

The BEARWATCH project depended on the cultural practices and traditional knowledge of the Inuit, and for that we are incredibly grateful. The support and collaboration of the Inuvialuit Game Council, the Gjoa Haven Hunters and Trappers Association, the Aiviit Hunters and Trappers Organization, the Canadian Rangers, & all the communities participating was integral to this project. The Governments of Nunavut and Northwest Territories provided key logistical and financial in-kind support for all aspects of sampling and we are grateful to them. This work was funded by the Government of Canada through Genome Canada and the Ontario Genomics Institute (OGI-123).

## Author Contributions

SNV conceived the study, conducted lab work and analyses, interpreted results, and wrote the first draft of the manuscript. SCL, M-JF and PVCdG conceived the study, interpreted results, and edited the manuscript. AGS interpreted results and edited the manuscript. RBGC-C, ELJ, and ZS conducted lab work. SCL, KM, MB, MD, and PVCdG, secured funding, built relationships with Inuit communities and organizations, managed samples, and reported results back to Inuit communities and Ontario Genomics and Genome Canada.

