## Supplementary methods, tables and figures for "Changes in sea ice alter genetic structure of an iconic Arctic apex predator in less than three decades"

[1] Department of Biology, Queen's University, Kingston, Ontario, Canada, [2] Department of Ecology and Evolutionary Biology, University of Toronto, Toronto, Ontario, Canada| [3] Hakai Institute, British Columbia, Canada, [4] School of Natural and Environmental Sciences, Newcastle University, Newcastle, UK.| [5] Department of Environment, Government of Nunavut, Igloolik, Nunavut, Canada, [6] Department of Environment and Natural Resources, Government of the Northwest Territories, Inuvik, Northwest Territories, Canada.

[†] deceased

### *Supplementary methods*

##### *S1. DNA extraction (adapted from Aljanabi and Martinez, 1997)*

Approximately 2-4 mm3 of tissue was subsampled from focal tissues using sterile surgical scissors and forceps, and added to a 1.5 mL Seal-Rite® microcentrifuge tubes (USA Scientific, Florida, USA) with 10 µL of proteinase K, 200 µL of sterile salt homogenizing buffer, 20 µL of 20% Sodium dodecyl sulfate (SDS) for digestion. Samples were vortexed and incubated at 56°C for at least 3 hours or until the tissues were fully lysed, with intermittent vortexing to increase lysis efficacy. PureLinkTM RNase A (Thermo Fisher Scientific, Waltham, Massachusetts, USA) treatment was added to each sample, which was then incubated at room temperature for 10 minutes, 150 µL of 5M NaCl added, and samples vortexed. I centrifuged the samples for 10 m at 12,000 rpm, and transferred the supernatant to 1.5 mL microcentrifuge tubes. 300 µL of cold (20°C) 100% ethanol was added and tubes inverted before incubating at -20°C for 1 hour or overnight. Samples were centrifuged at 4°C for 15 m at 12,000 rpm and supernatant discarded. 150 µL 70% ethanol was added and samples centrifuged at 4°C for 5 m, followed by discarding the supernatant and inverting the tubes to air-dry. Samples were then re-suspended in 80 µL of UltraPureTM DNase/RNase-Free Distilled Water (Thermo Fisher Scientific) and stored at -20°C.

***S2. DNA quality***

I evaluated DNA extract quality using 1.5% agarose gel electrophoresis stained with RedSafe® Nucleic Acid Staining solution (INtRON Biotechology, Gyeonggi-do, South Korea) against a 100 bp DNA ladder (Frogga Bio, Toronto, Ontario, Canada). Samples were subsequently evaluated for DNA quantity using a Nanodrop ND_1000 spectrophotometer (Thermo Fisher Scientific). Based on visual inspection of extracts and gel electrophoresis, when necessary, I used Solid Phase Reversible Immobilization (SPRI) magnetic beads, prepared using the Serapure protocol by Faircloth and Glenn (2011), to remove fat and cellular debris and purify DNA extracts. Magnetic beads were added to samples and incubated for 10 m. Samples were placed on a magnetic rack and separated for 2 m before discarding the supernatant. Beads were washed twice with 200 µL 70% ethanol and dried for 3-5 minutes before resuspension in 44 µL of UltraPure, and recovery of the solution. Samples were subsequently quantified using the Nanodrop ND_1000 spectrophotometer.

***S3. ddRADseq library preparation (adapted from Peterson et al., 2012)***

ddRADseq libraries were generated following a protocol developed by Peterson et al. (2012) and adjusted by Jensen et al. (2020). 1000 ng of extracted DNA was dispensed onto a plate and UltraPure was added up to 40 µL. One µL PstI-HF restriction enzyme, 1 µL MluCI restriction enzyme, 5 µL CutSmart® Buffer (New England Biolabs, Ipswich, Massachusetts, USA), and 3 µL UltraPure per reaction were aliquoted to each 40 µL DNA sample solution and incubated at 37°C in a thermocycler for 3 hours. I used SPRI magnetic beads to clean samples as per the protocol described above and 44 µL recovered. A QFX Fluorometer (DeNovix, Wilmington, Delaware, USA) with a proprietary high-sensitivity kit was used to quantify the samples prior to normalization. Ligation molarity calculations were used to normalize the concentration of DNA across individuals and 1.5 µL of annealed P1 and P2 adapters added in a combination particular to each individual 40 µL DNA sample solution. 5 µL NEB ligase buffer, 0.5 µL T4 ligase, and 1.5 µL ddH2O were added to each sample before incubation at 16°C for 90 m in the thermocycler, followed by denaturation at 65°C for 10 m, cooling for 90 s at 2°C and storage at 4°C. Ligation reactions were combined and subsequently redistributed, and SPRI bead clean-up was used twice more to clean the library pools, and resuspending in 60 µL and 30 µL of UltraPure, respectively. A BluePippin Prep (Sage Science, Beverly, Massachusetts, USA) was used according to the instructions and a size selection window of 400 to 490 base pairs (bp) with a 2% agarose cassette and 10 µg per lane. 3 µL of each library pool was amplified in eight parallel polymerase chain reactions (PCR) in an Eppendorf Mastercycler Gradient (Eppendorf, Hauppauge, New York, USA). Amplification in the thermocycler occurred by 98°C for 30 s followed by 10 cycles of: denaturation at 98°C for 10 s, primer annealing at 72°C for 30 s, and elongation at 72°C for 30 s, followed by a final elongation at 72°C 10 m, and cooling to 4°C. I combined PCR products and cleaned amplified pools with SPRI beads, resuspending these in 27 µL of UltraPure. Libraries were quantified using a QFX Fluorometer (DeNovix, Wilmington, Delaware, USA) before 200 pg of each barcoded library was sent to The Centre for Advanced Genomics (TCAG, Toronto, Ontario, Canada).

***S4. GT-seq library preparation (adapted from Campbell et al., 2015)***

GT-seq libraries were generated using a protocol developed by Campbell et al. (2015) and modified under BEARWATCH (Supplemental material; Hayward et al., 2021). Purified DNA was pipetted onto 96-well plates and treated with Exonuclease I (NEB M0293S: New England Biolabs, Ipswich, Massachusetts, USA)) and shrimp alkaline phosphatase (NEB M0731S) prior to the first PCR to remove single stranded DNA. PCR 1 amplification in the Eppendorf Mastercycler Gradient occurred by 95°C for 15 m followed by 5 cycles of: 95.0°C for 30s, 95.0°C for 3s , then -0.5°C per cycle at ~0.3°C/sec for 78 cycles until reaching 57.0°C, hybridization at 57.0° C for 30 s, and 72.0°C for 2 m followed by 15 cycles of specific amplification at 95.0°C for 30 s, 65.0°C for 30 s, and 72.0°C for 30 s and cooling to 4°C. We diluted PCR 1 products 20-fold with 133 µL of Nuclease Free H2O, tagged, and indexed these with 6-nucleotide primers, with unique barcodes on the 5' primer for each sample on a 96-well plate, and a unique barcode on the 7' primer for each plate, before PCR 2. PCR 2 in the Eppendorf Mastercycler Gradient occurred by 95.0°C for 15 m, followed by 10 cycles of 95.0°C for 10 s, 65.0°C for 30 s, and 72.0°C for 30 s, followed by 72.0°C for 5 m before cooling to 4.0°C. PCR 2 products were normalized with Just-a-Plate™ 96 PCR Purification and Normalization Kits (JN-120-10: Charm Biotech, Cape Girardeau, Missouri, USA), to elute 100 bp-20 kbp, 25-30 ng of DNA per sample. Tagged and indexed DNA was pooled, bead selected, and quantified using NEB library quantification kit (NEB E7630) following the protocol.

**S5.** ***Metaddata*****,** ***Genomic data demultiplexing and filtering***

We filtered individuals based on the usability of associated metadata and suitability for our analyses with VCFtools. Hunter recorded 'kill year' was used to infer the date each individual was sampled. When 'kill year' was unavailable, I used 'harvest year' instead. 'Harvest year', which at times was listed as the year following the 'kill year', may be biased due to killed bears being carried over to a later harvest period to abide by associated quotas for the area (for example, killings in defense or other non-tag kills). Hunter-assigned sex was recorded at the time of harvest, and these assignments were used for the purposes of sex-based analysis. As Circuitscape (McRae 2006) does not permit calculations for more than one individual per single focal node or grid cell, individuals were excluded based on individual missingness, retaining geographically coinciding individuals with more data.

We had sequence data from 1919 individuals from three independent sequencing batches: 544 individuals were successfully genotyped by ddRADseq and demultiplexed in January 2019, 415 individuals genotyped by GT-seq were demultiplexed in February 2020, and 960 individuals genotyped by GT-seq were demultiplexed in March 2021. 281 individuals were removed after filtering for minimum depth, genotype quality, and maximum 30% missingness across the 322 SNP panel. Forty-three technical replicates with greater missingness were removed. 381 fecal samples and 40 tissue set samples were removed due to poor metadata and duplications. 28 individuals were removed due to incomplete metadata. This left 1092 sequenced individuals with less than 30% missingness resulting from analyzing tissue-based samples, with unique locations and sufficient metadata from 1997–2020. Upon examining pairwise genetic distances between individuals, the individual missingness threshold was revised to a maximum acceptable threshold of 5%, yielding 867 individuals with acceptable data combined among the three sequencing batches, which are used in all subsequent analyses.

*Supplementary tables*

**Table S1**. Multi-surface (ice + landcover) resistance assignments of average proportional ice extents, averaged between 1997–2008 and 2009–2020 periods.

| Proportion ice | Reclassification | 1997–2008 MS_optim | 2009–2020 MS_optim | Mean (rounded to nearest integer) |
|---|---|---|---|---|
| Land | 1 | NA | NA | 0 |
| 1 | 2 | 2.06 | 274.64 | 138 |
| 0.9-0.999999 | 3 | 21.50 | 45.93 | 34 |
| 0.8-0.899999 | 4 | 1.71 | 22.58 | 12 |
| 0.7-0.799999 | 5 | 95.65 | 54.37 | 75 |
| 0.6-0.699999 | 6 | 201.82 | 65.72 | 134 |
| 0.5-0.599999 | 7 | 195.83 | 26.06 | 111 |
| 0.4-0.499999 | 8 | 403.78 | 218.88 | 311 |
| 0.3-0.399999 | 9 | 500.73 | 352.85 | 427 |
| 0.2-0.299999 | 10 | 493.70 | 458.56 | 476 |
| 0.1-0.199999 | 11 | 406.32 | 224.45 | 315 |
| 0-0.99999 | 12 | 440.85 | 462.49 | 452 |

**Table S2.** Categorical resistance results from single surface optimizations of 1997–2008 ice and landcover data using polar bear genetic distances from the Inuvialuit Settlement Region and Nunavut. Ice categories are measured in presence by proportion of each period. Resistances represent the relative resistance to movement of each category to gene flow.

| Surface | ice1997-2008 | | landcover | |
|---|---|---|---|---|
| | Feature | Resistance | Feature | Resistance |
| **Feature1** | *terrestrial* | 2 | *marine* | 2 |
| **Feature2** | 1 | 426 | temperate or sub-polar needleleaf forest | 259 |
| **Feature3** | 0.9-0.999999 | 108 | sub-polar taiga needleleaf forest | 652 |
| **Feature4** | 0.8-0.899999 | 1 | temperate or sub-polar broadleaf deciduous forest | 619 |
| **Feature5** | 0.7-0.799999 | 123 | mixed forest | 520 |
| **Feature6** | 0.6-0.699999 | 576 | temperate or sub-polar shrubland | 448 |
| **Feature7** | 0.5-0.599999 | 990 | temperate or sub-polar grassland | 936 |
| **Feature8** | 0.4-0.499999 | 1000 | sub-polar or polar shrubland-lichen-moss | 763 |
| **Feature9** | 0.3-0.399999 | 953 | sub-polar or polar grassland-lichen-moss | 138 |
| **Feature10** | 0.2-0.299999 | 685 | sub-polar or polar barren-lichen-moss | 87 |
| **Feature11** | 0.1-0.199999 | 286 | wetland | 270 |
| **Feature12** | 0-0.999999 | 619 | barren lands | 1 |
| **Feature13** | NA | NA | urban | 694 |
| **Feature14** | NA | NA | water | 902 |
| **Feature15** | NA | NA | snow and ice | 1000 |

**Table S3.** Categorical resistance results from single surface optimizations of 2009–2020 ice and landcover data using polar bear genetic distances from the Inuvialuit Settlement Region and Nunavut. Ice categories are measured in presence by proportion of each period. Resistances represent the relative resistance to movement of each category to gene flow.

| Surface | Ice2009-2020 | | landcover | |
|---|---|---|---|---|
| | Feature | Resistance | Feature | Resistance |
| **k** | | 13 | | 16 |
| **Feature1** | *terrestrial* | 1 | *marine* | 1 |
| **Feature2** | 1 | 740 | temperate or sub-polar needleleaf forest | 396 |
| **Feature3** | 0.9-0.999999 | 119 | sub-polar taiga needleleaf forest | 378 |
| **Feature4** | 0.8-0.899999 | 80 | temperate or sub-polar broadleaf deciduous forest | 397 |
| **Feature5** | 0.7-0.799999 | 124 | mixed forest | 782 |
| **Feature6** | 0.6-0.699999 | 144 | temperate or sub-polar shrubland | 562 |
| **Feature7** | 0.5-0.599999 | 96 | temperate or sub-polar grassland | 693 |
| **Feature8** | 0.4-0.499999 | 387 | sub-polar or polar shrubland-lichen-moss | 83 |
| **Feature9** | 0.3-0.399999 | 288 | sub-polar or polar grassland-lichen-moss | 21 |
| **Feature10** | 0.2-0.299999 | 675 | sub-polar or polar barren-lichen-moss | 44 |
| **Feature11** | 0.1-0.199999 | 213 | wetland | 56 |
| **Feature12** | 0-0.999999 | 329 | barren lands | 2 |
| **Feature13** | NA | NA | urban | 319 |
| **Feature14** | NA | NA | water | 32 |
| **Feature15** | NA | NA | snow and ice | 559 |

**Table S4.** Mantel R correlations at the first six distance lags for 1997–2008 (n = 126) and 20092020 (n = 126). Significant and interpretable correlations are bolded.

| | 1997–2008 | | | 2009–2020 | |
|---|---|---|---|---|---|
| **Lag (km)** | **Mantel R** | ***P*** | **Lag (km)** | **Mantel R** | ***P*** |
| 107.0 | **0.135** | **0.001** | 107.3 | **0.129** | **0.001** |
| 321.0 | **0.142** | **0.001** | 321.8 | **0.093** | **0.001** |
| 535.0 | **0.101** | **0.001** | 536.4 | **0.086** | **0.001** |
| 749.0 | **0.046** | **0.045** | 750.9 | 0.034 | 0.110 |
| 963.0 | -0.138 | 0.430 | 965.4 | -0.005 | 0.745 |
| 1177.0 | -0.043 | 0.013 | 1180.0 | -0.071 | 0.001 |

**Table S5.** Isolation-by-distance for each cluster identified in Figure 4c-f, between consistent sampling of time periods 1997–2008 and 2009–2020. Significant relationships are in boldface.

| | 1997–2008 | | | 2009–2020 | | |
|---|---|---|---|---|---|---|
| **Cluster** | ***n*** | **Mantel R** | ***P*** | ***n*** | **Mantel R** | ***P*** |
| Beaufort Sea | 15 | 0.15 | 0.17 | 15 | 0.001 | 0.989 |
| Arctic Archipelago | 89 | **0.186** | **0.002** | 89 | **0.138** | **0.01** |
| Hudson Complex | 22 | **0.17** | **0.041** | 22 | 0.14 | 0.077 |

**Table S6.** Isolation-by-distance between sexes and overall for 1997–2008 and 2009–2020. Significant relationships are in boldface.

| | 1997–2008 | | | 2009–2020 | | |
|---|---|---|---|---|---|---|
| **Subset** | ***n*** | **Mantel R** | ***P*** | ***n*** | **Mantel R** | ***P*** |
| Females | 199 | **0.31** | **0.001** | 173 | **0.29** | **0.001** |
| Males | 247 | **0.25** | **0.001** | 248 | **0.28** | **0.001** |
| Overall | 446 | **0.28** | **0.001** | 421 | **0.28** | **0.001** |

**Table S7.** sPCA Monte-Carlo tests based on 499 permutations suggest the presence of at least one significant global structure and no significant local structure for either period.

| 1997-2008 spca_randtest() | 2009-2020 spca_randtest() |
|---|---|
| Global Monte-Carlo test<br>Observation: 5.14<br>*P* = 0.002 | Global Monte-Carlo test<br>Observation: 5.54<br>*P* = 0.002 |
| Local Monte-Carlo test<br>Observation: 4.16<br>*P* = 0.992 | Local Monte-Carlo test<br>Observation: 4.42<br>*P* = 1.0 |

**Table S8**. Summary table of 1997–2008 bootstrap analysis results following 1000 iterations. %.top indicates the frequency that each model was identified as the top model.

| Surface | av.AIC | av.AICc | av.weight | av.rank | n | %.top | k |
|---|---|---|---|---|---|---|---|
| **distance** | 10616.8 | 10617.0 | 0.999 | 1 | 1000 | 100 | 2 |
| **ice** | 10637.8 | 10642.4 | < 0.001 | 2.003 | 0 | 0 | 13 |
| **landcover** | 10644.6 | 10651.7 | < 0.001 | 2.998 | 0 | 0 | 16 |
| **landcover + ice1997_2008** | 10661.6 | 10686.6 | < 0.001 | 3.999 | 0 | 0 | 28 |

**Table S9.** Summary of 2009–2020 bootstrap analysis results following 1000 iterations. %.top indicates the frequency that each model was identified as the top model.

| Surface | av.AIC | av.AICc | av.weight | av.rank | n | %.top | k |
|---|---|---|---|---|---|---|---|
| **landcover + ice2009_2020** | 10488.2 | 10513.1 | 0.854 | 1.02 | 979 | 97.9 | 28 |
| **distance** | 10542.4 | 10542.5 | 0.133 | 2.43 | 19 | 1.9 | 2 |
| **landcover** | 10544.3 | 10551.4 | 0.0135 | 2.55 | 2 | 0.2 | 16 |
| **ice2009_2020** | 10939.0 | 10943.5 | < 0.001 | 4 | 0 | 0 | 13 |

**Table S10.** Generalized least squares models fitted with a Gaussian spatial correlation structure to predict polar bear genetic change (GC) across the Canadian Arctic using two different metrics of landscape change. RES – change in optimized resistance between periods, averaged between sea ice parameterizations for either period; DEG – direction of sea ice change in either positive or negative degrees of change.

| Model (GC ~) | AIC | ΔAIC | wAIC |
|---|---|---|---|
| $\beta_{DEG} + \beta_{RES}$ | 51889.8 | 0 | 1.0 |
| $\beta_{DEG}$ | 51901.1 | 11.2 | 0 |
| $\beta_{RES}$ | 51915.8 | 25.9 | 0 |

*Supplementary figures*

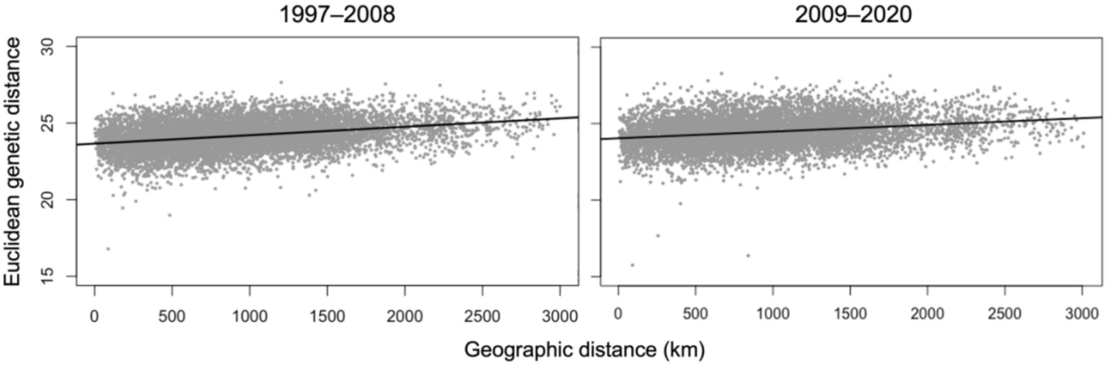


**Figure S1.** Polar bear isolation-by-distance plots for 1997–2008 (R = 0.32, P = 0.001, n = 126) and 2009–2020 (R = 0.24, P = 0.001, n = 126) subsets of consistent sampling across Nunavut and the Inuvialuit Settlement Region.

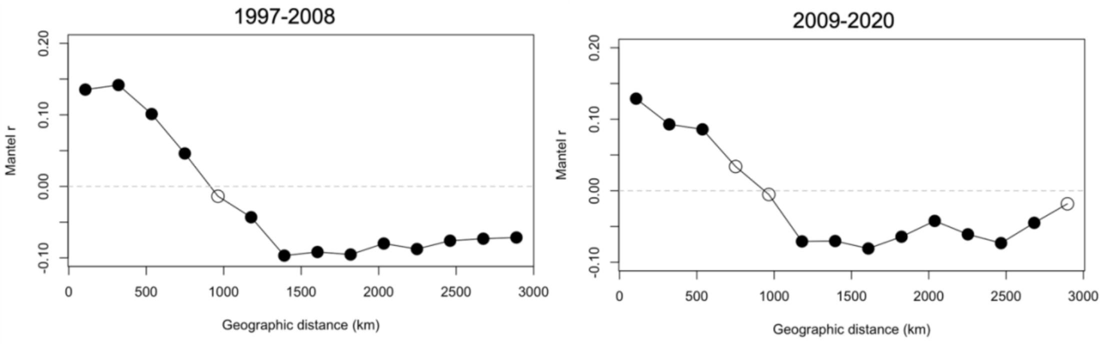


**Figure S2.** Mantel correlograms for 1997–2008 (n = 126) and 2009–2020 (n = 126) over distance lags defined by Sturges Rule.

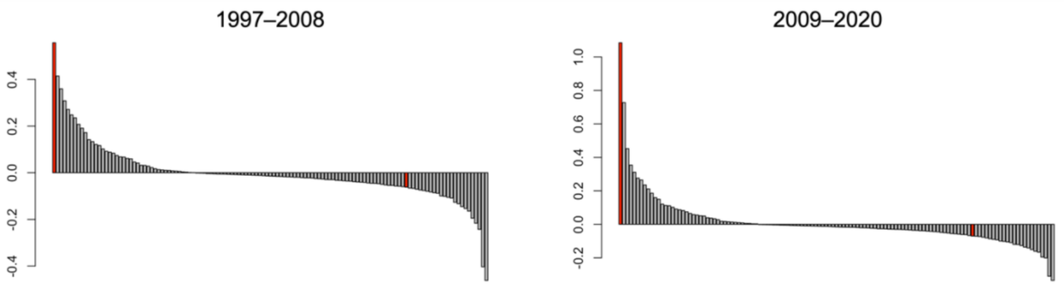


**Figure S3.** Barplot of eigenvalues 1997–2008 and 2009–2020 spatial analyses of principal components. 1st and 100th eigenvalues are indicated in red.

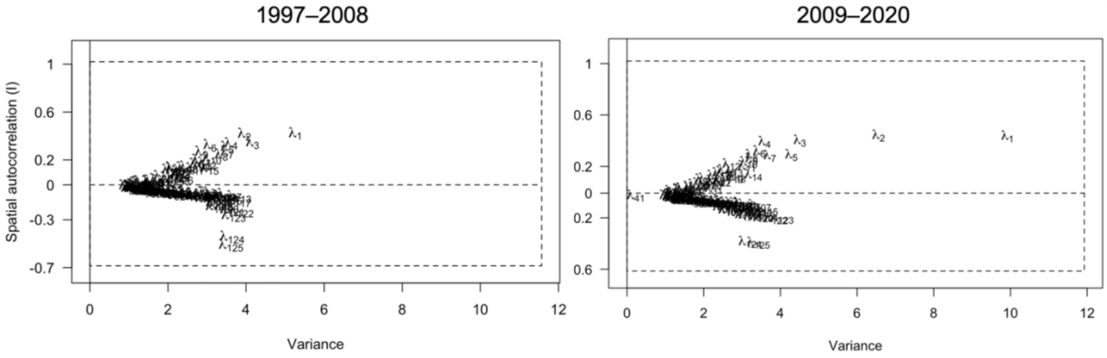


**Figure S4.** Screeplots of genetic variance and spatial autocorrelation for 1997–2008 and 20092020 spatial analyses of principal components.
